# Investigating the human and non-obese diabetic mouse MHC class II immunopeptidome using protein language modelling

**DOI:** 10.1101/2022.08.19.504560

**Authors:** Philip Hartout, Bojana Počuča, Celia Méndez-García, Christian Schleberger

## Abstract

Identifying peptides associated with the major histocompability complex class II (MHCII) is a central task in the evaluation of the immunoregulatory function of therapeutics and drug prototypes. MHCII-peptide presentation prediction has multiple biopharmaceutical applications, including the safety assessment of biologics and engineered derivatives *in silico*, or the fast progression of antigen-specific immunomodulatory drug discovery programs in immune disease and cancer. This has resulted in the collection of large-scale data sets on adaptive immune receptor antigenic responses and MHC-associated peptide proteomics. In parallel, recent deep learning algorithmic advances in protein language modelling (PLM) have shown potential in leveraging large collections of sequence data and improve MHC presentation prediction. Here, we train a compact transformer model (AEGIS) on human and mouse MHCII immunopeptidome data, including a preclinical murine model, and evaluate its performance on the peptide presentation prediction task. We show that the transformer performs on par with existing deep learning algorithms and that combining datasets from multiple organisms increases model performance. We trained variants of the model with and without MHCII information. In both alternatives, the inclusion of peptides presented by the I-A^g7^ MHC class II molecule expressed by the non-obese diabetic (NOD) mice enabled for the first time the accurate*in silico* prediction of presented peptides in a preclinical type 1 diabetes model organism, which has promising therapeutic applications.

**Availability and implementation:** The source code is available at https://github.com/Novartis/AEGIS.

## 1 Introduction

The major histocompatibility complex class II (MHCII) is expressed on the surface of professional antigen presenting cells (APCs). Its main function is the presentation of peptides from extracellular proteins to subpopulations of CD4^+^ T cells (Wieczorek et al., 2017). Co-stimulatory molecules expressed upon recognition influence CD4 ^+^ T cell activation and differentiation into either regulatory (T _reg_) or helper T cells (T_h_), which locally induce tolerance or promote inflammation (Mueller, 2010; Pepper and Jenkins, 2011; Graham and Xavier, 2020; Murphy and Murphy, 2021). Peptide presentation prediction by the MHCII has numerous biopharmaceutical use cases, such as enabling rapid progress in the development of immunomodulatory therapeutic strategies, or assisting in the selection of non–immunogenic biologic drugs. Furthermore, computational selection of peptides for antigen–specific immune modulation deepens our mechanistic understanding in autoimmune disease and immune escape in cancer (Zhang et al., 2021).

The MHCII is a heterodimer containing a b and an a chain, encoded by the human leukocyte antigen (HLA) gene complex. The structural features of the molecule influence peptide presentation. For instance, the binding groove ends of the MHCII are open, which allows for a higher diversity of ligands to be presented, compared to the closed-end groove of the MHC class I (Laimer and Lackner, 2021; Wu et al., 2021). This promiscuity poses a major challenge for *in silico* techniques because long stretches of peptides do not contribute to the binding core, while the edges of the peptide could still contain information regarding the digestion process in the multivesicular antigen-processing intracellular compartments of the APC (Reinherz et al., 1999; Schneidman-Duhovny et al., 2018). Bioinformatic methods dedicated to the prediction of peptides presented by the MHCII need to take into account the binding core, the fraction of the peptide that interacts with the binding pocket of the MHCII molecule, and the amino acids that contribute the most to the binding core, referred to as anchor residues.

There are multiple strategies leveraging binding cores and anchor amino acids for the presentation prediction task, which is necessary for the classification of presented and non-presented (or seldom-presented) peptides. These include position-specific scoring matrices (Nielsen et al., 2008), standard perceptron-based neural networks (Reynisson et al., 2020), convolutional neural networks (Racle et al., 2019), or transformers (Cheng et al., 2021). However, the variety of peptides presented poses significant challenges, lowering the performance of *in silico* methods. In this work, we aimed to leverage transformers due to their ability to capture long and varying ranges of dependencies in data, their computational efficiency, and their capability to handle input data of different dimensions.

We apply recent advances in natural language processing algorithms to peptides in order to investigate their advantage in the identification of the MHCII immunopeptidome in humans and a preclinincal mouse model. We evaluated a transformer encoder on a predominantly human peptide:MHCII dataset and a second containing exclusively pancreatic islet MHCII peptidome data of non-obese diabetic (NOD) mice to assess predictive potential tailored to specific project needs. We validate the relative performance of our model against others published in the literature on the same data and determine that it outperforms previously published strategies on the available feature sets.

## 2 Methods

### 2.1 Datasets

To build the model, we sourced human (H) and mouse (M) peptide data from MHCII binding affinity (BA) and mass spectrometry (MS)-eluted ligand (EL) assays from the immune epitope database (IEDB) (Vita et al., 2019). The EL subset contains exclusively single-allele (SA) peptide:MHCII information for nearly 1000 human (HLA-DR, HLA-DQ, and HLA-DP) and wild type mouse (H–2) alleles. These data were utilized together with multiallele (MA) information by Reynisson et al. (2020) to train NetMHCIIpan v4.0. The BA data includes peptide:MHCII binding affinities retrieved from the IEDB and partially overlaps with the dataset used to generate NetMHCII v2.3 and NetMHCIIpan v3.2 by Jensen et al. (2018). A total of 447232 peptides associated to 85 unique alleles were represented in the BA and EL IEDB-sourced datasets. These experimentally-acquired (presented) peptides were distributed into folds as follows: 90-95% of the data was assigned to the training set, 2.5-5% to the validation set, and 2.5-5% to the test set (Table 1).

**Table 1:** Descriptive statistics of the IEDB and NOD I-A^g7^ datasets used for training, validation, and evaluation of the model. IEDB contains both human (H) and mouse (M) peptides from BA and EL assays. The NOD I-A^g7^ repository consists of EL data only. ^*^: No decoy BA data was generated (see section 2.2 for further explanation). The distribution of the BA data labels can be found on Supplementary Figure 1. ^**^: These allele totals represent the total amount of unique alleles.

| Data Source<br>(Organism) | Data type<br>(n alleles) | Experimentally<br>acquired |  |  | Synthetic |  |  |
| --- | --- | --- | --- | --- | --- | --- | --- |
|  |  | Train | Validation | Test | Train | Validation | Test |
| IEDB (H) | BA<sup>*</sup>(n=65) | 97223 | 4834 | 6109 | — | — | — |
|  | EL (n=19) | 286010 | 6965 | 7155 | 2469500 | 64950 | 64920 |
| **Total** | 77<sup>**</sup> | 383233 | 11799 | 13264 | 2469500 | 64950 | 64920 |
| IEDB (M) | BA<sup>*</sup>(n=8) | 2308 | 30 | 68 | — | — | — |
|  | EL (n=2) | 35085 | 695 | 750 | 385195 | 10055 | 10055 |
| **Total** | 8<sup>**</sup> | 37393 | 725 | 818 | 385195 | 10055 | 10055 |
| NOD I-A<sup>g7</sup> (M) | EL (n=1) | 4669 | 467 | 258 | 23345 | 2335 | 1290 |

We set out to evaluate the usability of limited project–specific information by considering MHCII peptidome data from NOD mice, which express the I-A^g7^ variant of the MHCII molecule. We used the dataset generated by Wan et al. (2020), which includes peptides eluted from myeloid APCs contained in pancreatic islets, peripheral lymph nodes, and spleens of NOD mice. A total of 5394 EL-only data points were split into folds as described above (Table 1).

### 2.2 Negative data (decoys)

The nature of the EL data acquisition workflow does not allow for the identification of non-presented peptides. These are required to train the binary machine learning classification model and were, therefore, generated *in silico*. The synthetic (negative data, decoys, non-presented, rarely-presented) peptides generated by Reynisson et al. (2020) were utilized. To create this set, peptides from the UniProt database were randomly and uniformly sampled for each length between 15 and 21 amino acids, including peptide flanking regions (PFRs) (UniProt Consortium, 2021). A more specific approach was adopted for the decoys derived from NOD mice, where random peptides were sampled from non-presented regions of proteins from which at least one peptide was presented. This ensured that relevant decoys were included in the model. For BA data, the experiment identifies rarely-presented as well as non-presented peptides. The affinity data points were log-transformed to a scale between 0 and 1 from experimental IC_50_ binding values as follows: 1–log(IC_50_ nM)/log (50 000) (Nielsen et al., 2003; Jensen et al., 2018) (Supplementary Figure 1).

### 2.3 Data preparation

We relied on the method employed by NetMHCIIpan v3.0 to represent the MHCII binding pocket (Karosiene et al., 2013). In brief, a subset of amino acids was selected according to crystal structures of peptide:MHCII complexes. The residues considered to be part of the binding pocket are located within 4.0 Å of the presented peptide. The resulting positions were used to build a pseudo sequence of the HLA, containing the most important amino acids determining the specificity of the binding pocket. It is worth noting that most of the polymorphism of the MHC regions occurred at those positions, further supporting the use of this feature compression method for the HLA. We extracted the pseudo sequence of I-A^g7^ by aligning the structure of I-A^g7^ (PDB-ID 1f3j) to HLA-DR3 (PDB-ID 1a6a) and identifying the corresponding residues in I-A^g7^, as described in Karosiene et al. (2013). (Supplementary Figure 2).

The resulting feature set included the raw eluted peptide, ranging between 9-15 amino acids long, and the pseudo sequence. In addition, we built model variants without MHCII pseudo sequences to obtain a general presentability score regardless of MHCII allele.

An overview of the entire data processing pipeline can be found in Supplementary Figure 3.

### 2.4 External datasets

We used independently acquired external databases to benchmark the performance of our model against published methods. We focused on the data used to build the MARIA multimodal recurrent neural network (RNN) (Chen et al., 2019). These data include two ligandomes from K562 cells (leukemia-derived lymphoblasts) expressing the b-chain alleles HLA-DRB1^*^01:01 and HLA-DRB1*04:04, as well as an independent melanoma HLA-II ligand set. These data were chosen based on availability and comparability with the performance of different models. Unlike MARIA, our model does not integrate gene expression data, as this type of information is normally lacking. Additionally, expression data can only be obtained for endogenous proteins or peptide tiles (if a library is being evaluated), therefore excluding exogenous antibodies, peptides, or engineered derivatives assayed *in vitro*.

### 2.5 Modelling setup

A variant of the model was trained excluding pseudo sequence information for both IEDB and the NOD mice dataset, which poses an advantage when allele-specific information is not available.

To divide our data into training, validation, and test sets, we randomly split the peptides and MHCII combinations across folds and subsequently isolated overlapping peptides to ensure the datasets were independent. The pan-specificity (i.e., ability to achieve high accuracy in the classification task for unseen alleles) of some predictors is often touted in the literature. We maintain that utilizing the model on novel alleles is not a predominant use case and that learning patterns of presented peptides related to known MHCII molecules is more useful and accurate.

The architecture of the transformer model used in this study is shown in Figure 1. In brief, the sequences are first marked with start and end of sequence tokens, then padded to ensure consistency of input to the model while avoiding spurious predictive power extracted from the padded characters. Then, the sequences are fed through an embedding layer and a positional encoder to ensure that the position of each amino acid relative to one another is encoded into the embedding. Subsequently, the sequence of (i) multi-head attention networks, (ii) an addition and normalization layer, (iii) a feed-forward network, and (iv) one last addition and normalization layer are added and repeated four times. Prior to computing the final sigmoid value used for classification, one last feedforward step processes the output of the aforementioned block. We used a variety of well-established metrics to assess the binary classification performance of machine learning models on imbalanced datasets, such as the receiver operating characteristic (ROC) curve, as implemented through `scikit-learn’s metrics` API (Pedregosa et al., 2011).

**Figure 1:**
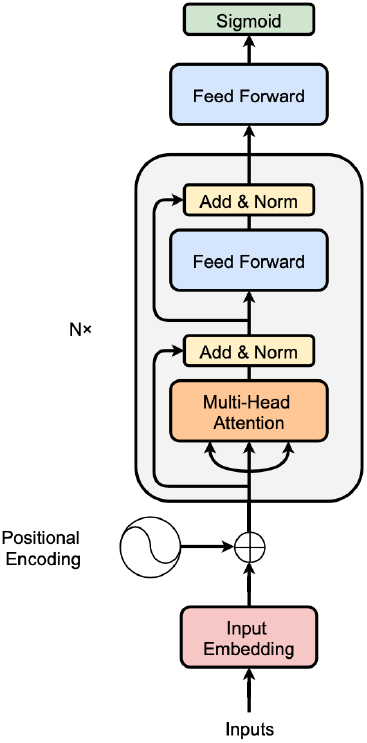
Transformer encoder architecture used to predict peptide presentation by MHCII; adapted from Vaswani et al. (2017). Instead of processing the data sequentially, the positional encoder embeds position-related information, which serves as input to a set of transformer blocks that extract features used for a small feedforward neural network, using these embeddings as features of prediction.

The transformer was trained using the binary cross-entropy loss function together with the Adam optimization algorithm for 200 epochs (Kingma and Ba, 2014). The model hyperparameters can be found in the code, available at https://github.com/Novartis/AEGIS. In total, the transformer has approximately 3.5M trainable parameters. We trained the model on 3 Tesla V100 GPUs.

Particular attention was placed into maintaining a well-documented modular pipeline, using best software engineering practices.

## 3 Results

The sample sizes of each of the datasets are shown in Table 1. Table 2 contains the performance of the HLA-agnostic model (i.e., no pseudo sequence is given as input, referred to as *Peptide only*) and the HLA-specific model, *Peptide:MHC*, both trained on IEDB, NOD, and IEDB + NOD data sources.

**Table 2:** Performance results averaged from five different runs for each of the five datasets.The values in the table represent the results for each setting’s respective validation set. The model performance was remarkably stable, with standard deviation values only affecting the performance on the fourth digit after the comma (± 0.0002 for all model configurations). ^*^ The pseudo sequence for this model is the same for all samples.

| Data source | Feature set | Metric |  |  |
| --- | --- | --- | --- | --- |
|  |  | AUC | F1 | MCC |
| IEDB | Peptide only | 0.935 | 0.671 | 0.646 |
|  | Peptide:MHC | 0.959 | 0.770 | 0.753 |
| NOD | Peptide only* | 0.775 | 0.185 | 0.164 |
| IEDB + NOD | Peptide only | 0.926 | 0.648 | 0.622 |
|  | Peptide:MHC | 0.955 | 0.756 | 0.736 |

The best performing models displayed an AUC > 0.95 and corresponded to the IEDB and IEDB + NOD independent and identically distributed (i.i.d.) datasets including psedosequence information (Peptide:MHC). These models underperformed considering metrics with set thresholds, such as the F1, accuracy, precision, and recall. While performance is on average lower for the mouse data, possibly due to overfitting, it is still usable, especially in use cases where a model with high recall (specificity) is required.

All four models were evaluated on the I-A^g7^ test set, to which the model had not been previously exposed (Table 3). The best performance (AUC=0.86) was newly observed for the i.i.d. IEDB + NOD dataset containing pseudo sequence information.

**Table 3:** Inference results from the top four models (NOD mouse-only model is excluded) evaluated on the test set that contains NOD mouse-only data (1630 peptides).

| Data source | Feature set | Metric |  |  |
| --- | --- | --- | --- | --- |
|  |  | AUC | F1 | MCC |
| IEDB | Peptide only | 0.55 | 0.1 | -0.11 |
|  | Peptide:MHC | 0.54 | 0.14 | -0.03 |
| IEDB + NOD | Peptide only | 0.8 | 0.19 | 0.14 |
|  | Peptide:MHC | 0.86 | 0.55 | 0.5 |

The model displayed a median AUC > 0.90 for the most frequent alleles of the HLA-DR, HLA-DQ, and HLA-DP human isotypes (Greenbaum et al., 2011), with best values observed for HLA-DRB1*15:01 (AUC=0.97), HLA-DRB1*16:01 (AUC=0.97), HLA-DRB1*07:01 (AUC=0.96), HLA-DRB1*01:01 (AUC=0.96), HLA-DRB3*01:01 (AUC=0.96), HLA-DRB1*04:05 (AUC=0.95), HLA-DPA1*01:03/DPB1*06:01 (AUC=0.95), HLA-DRB1*11:01 (AUC=0.95), HLA-DRB1*03:01 (AUC=0.95), HLA-DPA1*01:03/DPB1*02:01 (AUC=0.94), HLA-DQA1*01:02/DQB1*06:02 (AUC=0.93), HLA-DRB1*04:01 (AUC=0.92), and HLA-DRB1*12:01 (AUC=0.91). See Supplementary Tables 1-4 for further details.

The learning curves for each of the models can be found in Supplementary Figure 4.

### 3.1 Model performance on independent datasets

On all datasets, our model outperformed netMHCIIpan v4.0 and v3.2 by a substantial margin (HLA-DRB1^*^01:01: our model AUC=0.82, netMHCIIpan AUC=0.608; HLA-DRB1^*^04:04: our model AUC=0.82, netMHCIIpan AUC=0.568). It performs on par with the MARIA multimodal RNN, despite unavailability of gene expression data (HLA-DRB1^*^01:01: our model AUC=0.82, MARIA AUC=0.885; HLA-DRB1^*^04:04: our model AUC=0.82, MARIA AUC=0.892) (Figure 2).

**Figure 2:**
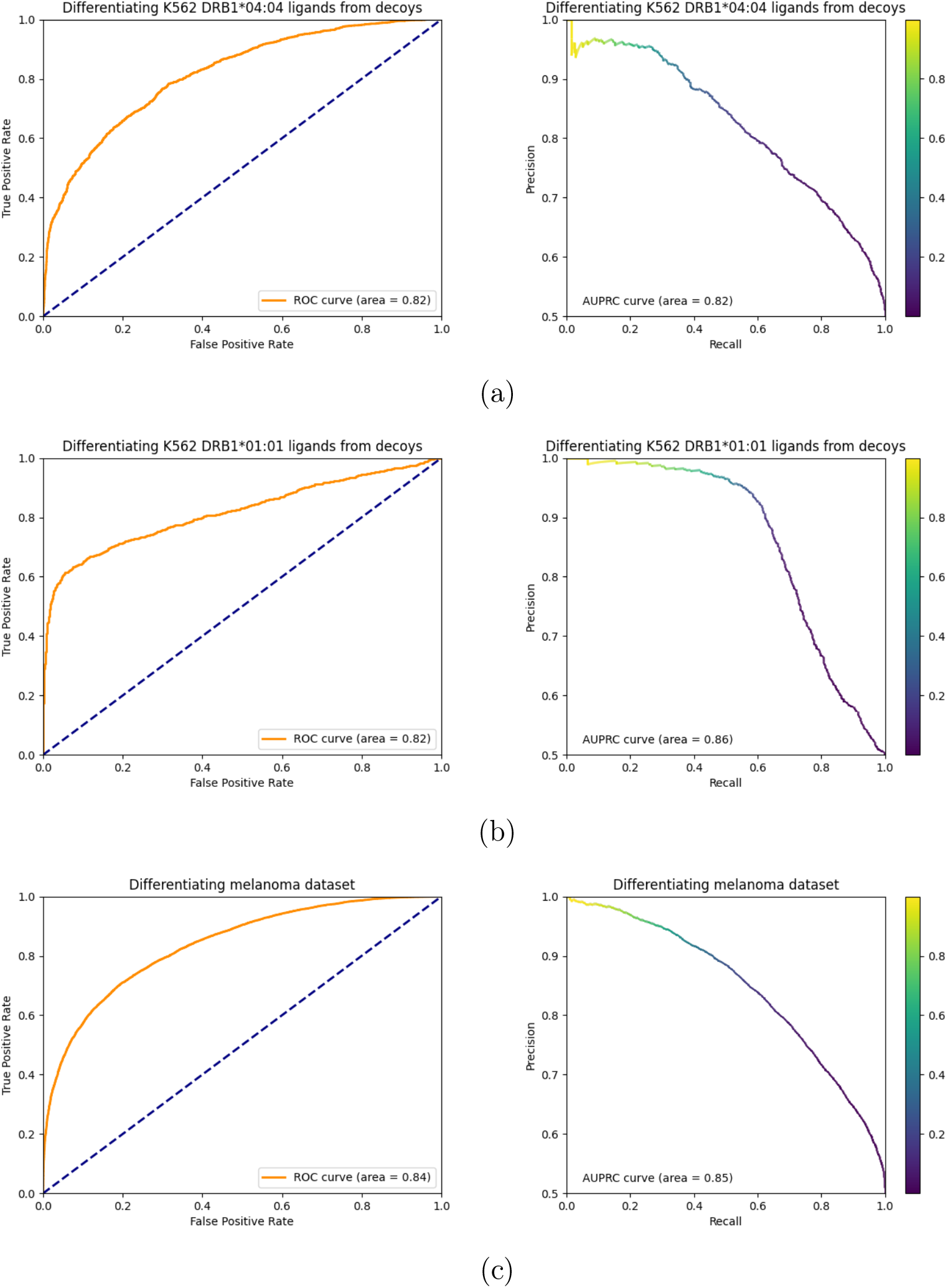
Receiver operating characteristic (ROC) and precision-recall curves of the transformer-based model on external datasets. Allele-specific model on the K652 cell-line data for the DRB1*04:04 (a) and DRB1*01:01 (b) alleles. (c) Performance of the allele-agnostic model on the melanoma dataset. The color gradient on the precision-recall curve indicates the operating threshold chosen to get the respective precision and recall value.

## 4 Discussion

Overall, our results show that protein language models hold promise in the prediction of peptide presentation by the MHCII. Specifically on independently acquired experimental data, the model presented here performs on par with models that leverage other features, such as gene expression. This is significant for biopharmaceutical settings where gene expression data cannot be acquired.

Our model performs comparably to BERTMHC, the first and most recent evaluation of a transformer neural network model on MHCII peptide data (Cheng et al., 2021). BERTMHC relies on the pretrained protein BERT (Bidirectional Encoder Representations from Transformers) model (Devlin et al., 2019). In brief, BERT is a bidirectionally–trained language model that learns the context of a word based on its surroundings, therefore enabling the extraction of a deeper sense of context than single-direction models. Computationally, BERT consists of 12 layers with 12 attention heads per layer. In contrast, our model consists of 4 layers with 2 attention heads per layer. Both transformers are trained on IEDB-sourced immunopeptidome data, also utilized to train NetMHCIIpan versions 3.2 (BA information) and 4.0 (extending to EL data), with slight differences in data sizes (see Cheng et al. (2021) and Table 1 for details). BERTMHC considers both multiallele (MA) and single-allele (SA) information, although a model trained in SA-only data (BERTMHC-SA) is also presented by Cheng et al. (2021). The performance of our model (SA data exclusively) is comparable to both BERTMHC and BERTMHC-SA, in spite of model size differences. In addition, both models are able to perform well (AUC > 0.75) on independent sets of data, consisting of patient–derived cancer immunopeptidome information in BERTMHC and two ligandomes from K562 cells and an independent melanoma MHCII ligand set reported by Chen et al. (2019) in our work.

We also describe an improved I-A^g7^-specific predictor which, despite leveraging a modest dataset for training, still performs satisfactorily on most metrics. The advantage of using such a transformer model on mouse data is that the training can be carried out in human, wild type, and NOD model mouse information simultaneously, hence applying patterns of immune recognition seen in other alleles and organisms while still identifying NOD mouse-specific signals. Thus far, the only available model for prediction of the I-A^g7^ immunopeptidome is PRED^NOD^, a scoring matrix–based method published by Rajapakse et al. (2006). The training and testing data of PRED^NOD^ were derived primarily from heterogeneous BA experiments, with a total of 653 experimentally-acquired binders and 1000 non–binders, whereas exclusively EL data (>20K peptides in the training set only) from a single experimental setup was utilized in our model. Our contribution offers a working alternative and extends the functionality of PRED^NOD^, suitable for the prediction of I-A^g7^ binders, rather than presented peptides.

While some important machine learning-related methodological advancements are presented here, there are still several sources of bias to consider and some shortcomings that need to be addressed in the future. First, to ensure feature consistency in our dataset, we omitted PFRs in our models. PFRs are the three residue positions before and after the peptide presented by the MHCII molecule on the source protein sequence. They have been reported to boost predictive performance and should be considered in future research (Barra et al., 2018).

There are also biases inherent to the data. We know that peptides with certain physicochemical properties allowing for good detection are over-represented in mass spectrometry analytics (Jarnuczak et al., 2016), whereas e.g., cysteine-containing peptides are under-represented in MS–generated datasets (Abelin et al., 2017). In addition, BA data selects for peptides that bind to the MHCII molecule, i.e., it exhibits biochemical signature matches to the MHCII molecules that may not be actually presented by the MHCII in nature, either because the peptide does not occur naturally or because it is being filtered by the complex intracellular dynamics involved in loading peptides onto the MHCII molecule (Karle, 2020). The addition of these type of data to the training set may reduce performance in the *in silico* assessment of drug immunogenicity, while still producing usable results.

Furthermore, while sampling from UniProt to generate synthetic negative peptides as done by Reynisson et al. (2020) is an accepted practice across the literature to generate synthetic counterparts to the positive MS-data, it is not experimentally validated, and may produce sequences that could be in fact presented by the MHCII molecules.

Additional shortcomings are the inability of our method to handle multi-allele data for model variants requiring a pseudo sequence of the MHCII as input. This would require a semi-supervised learning setting, where peptides eluted from multi-allele cultures are assigned the most likely MHCII that presented each of them. This multi-label classification task could be enabled by a first classifier that learns to assign one or more MHCII molecules to each of the eluted peptides resolved using MS. Once the labeling of the multi-allele dataset is achieved, it could then be used as a larger pool of peptides for the main classifier predicting peptide presentation. This approach has led to one of the leading techniques for peptide presentation to leverage multi-allele data and thus, a wider coverage of many more real-life scenarios (Nielsen and Lund, 2009; Reynisson et al., 2020). With this expanded dataset, more features could be added, such as information related to the peptide position in the protein structure and the source protein localization in the cell (Ten Broeke et al., 2013; Stern and Santambrogio, 2016). In addition, building a differentiable encoding of the binding pocket from the entire MHCII pseudo sequence, possibly using protein language modelling, could also prove to extract more informative features than the currently used pseudo sequence.

Finally, while the randomly sampled peptides from UniProt contain peptides originating from multiple organisms, expanding the repertoire of presented peptides to bacterial species could further enhance the potential of our model to microbiome-related applications. This has been investigated before (Graham et al., 2018), but protein language models could further enhance the predictive performance of such models. Additionally, we have shown that predictive performance in humans is sustained after the incorporation of MHCII equivalents from other organisms and that combining human data with information from different alleles and organisms might enable peptide presentation prediction by the MHCII molecule in species for which limited data is available.

## 5 Conclusion

Our work demonstrates that protein language models hold promise to tackle the difficult problem of identifying which peptides are presented by different MHCII alleles in homozygous data. Attention mechanisms are potent modelling strategies to identify short- and long-term dependencies between amino acids, which is ideal to extract features related to the PFRs, the binding core, and its interaction with the most important residues to determine the specificity of the open MHCII binding pocket. Additionally, we have demonstrated the first protein language model specifically trained on I-A^g7^ data. As such, it can support efforts directed at leveraging NOD mice immunopeptidome measurements and drive therapeutic innovation.

## Acknowledgements

We thank Sylvie Le Gall, Anette Karle, Michael Gutknecht, Barbara Brannetti, Jiayi Cox, Stephan Reiling, and Leona Gabryöová for the valuable scientific discussions.

## Financial Support

none declared.

## Conflict of Interest

P.H. and B.P. were temporarily employed by the Novartis Institutes for Biomedical Research (NIBR) while conducting the presented research. C.M-G. and C.S. are permanently employed by NIBR.

## Data availability

The data are archived permanently on Zenodo http://doi.org/10.5281/zenodo.7247911.

## Supplementary Information

### Supplementary tables

**Supplementary Table 1:** Model evaluation for an unbiased estimate of the model’s performance in the most frequent HLA-DRB1 alleles.

| HLA-DRB1<br>Metric | 01:01 | 03:01 | 04:01 | 04:05 | 07:01 | 08:02 | 09:01 | 11:01 | 12:01 | 15:01 | 16:01 |
| --- | --- | --- | --- | --- | --- | --- | --- | --- | --- | --- | --- |
| **AUC** | 0.96 | 0.95 | 0.92 | 0.95 | 0.96 | 0.72 | 0.77 | 0.95 | 0.91 | 0.97 | 0.97 |
| **F1** | 0.79 | 0.73 | 0.67 | 0.68 | 0.76 | 0.47 | 0.56 | 0.73 | 0.62 | 0.80 | 0.83 |
| **MCC** | 0.78 | 0.70 | 0.65 | 0.65 | 0.74 | 0.31 | 0.37 | 0.71 | 0.58 | 0.78 | 0.82 |

**Supplementary Table 2:**
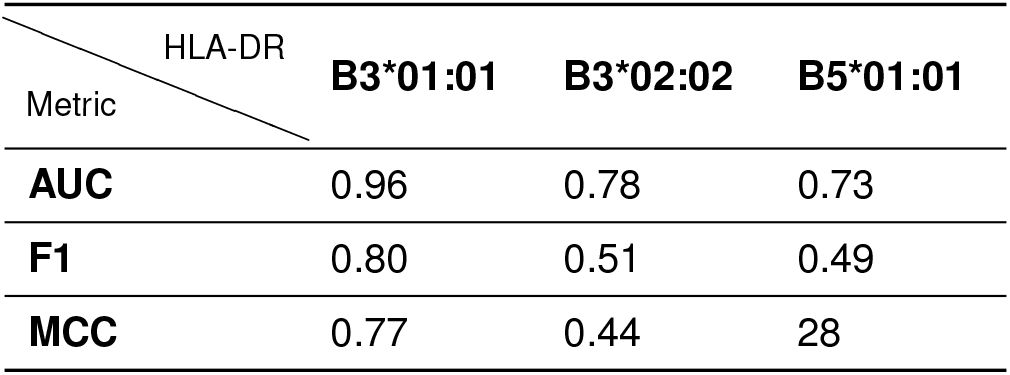
Model evaluation for an unbiased estimate of the model’s performance in additional most frequent HLA-DR alleles.

| HLA-DR<br>Metric | B3*01:01 | B3*02:02 | B5*01:01 |
| --- | --- | --- | --- |
| **AUC** | 0.96 | 0.78 | 0.73 |
| **F1** | 0.80 | 0.51 | 0.49 |
| **MCC** | 0.77 | 0.44 | 28 |

**Supplementary Table 3:**
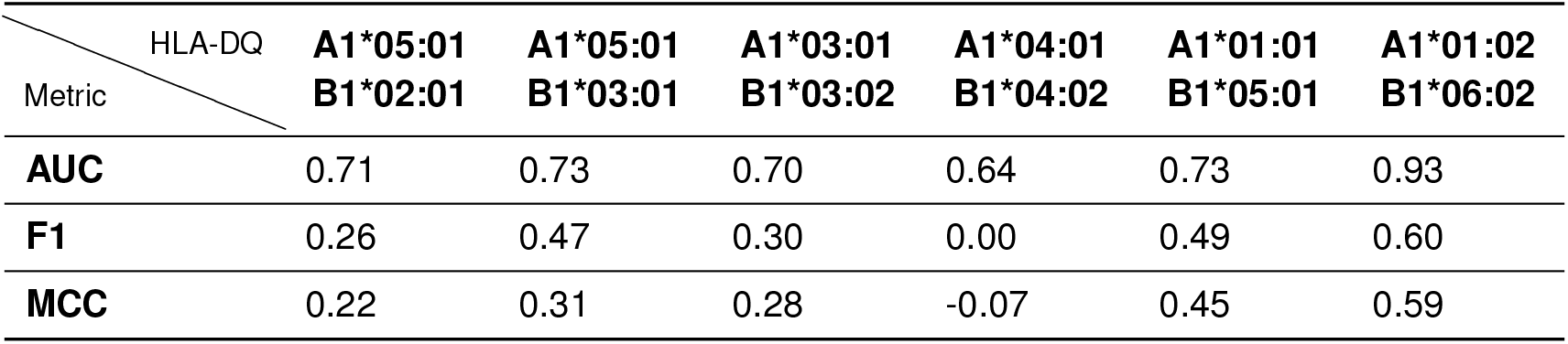
Model evaluation for an unbiased estimate of the model’s performance in the most frequent HLA-DQ allele pairs.

| HLA-DQ<br>Metric | A1*05:01<br>B1*02:01 | A1*05:01<br>B1*03:01 | A1*03:01<br>B1*03:02 | A1*04:01<br>B1*04:02 | A1*01:01<br>B1*05:01 | A1*01:02<br>B1*06:02 |
| --- | --- | --- | --- | --- | --- | --- |
| **AUC** | 0.71 | 0.73 | 0.70 | 0.64 | 0.73 | 0.93 |
| **F1** | 0.26 | 0.47 | 0.30 | 0.00 | 0.49 | 0.60 |
| **MCC** | 0.22 | 0.31 | 0.28 | -0.07 | 0.45 | 0.59 |

**Supplementary Table 4:** Model evaluation for an unbiased estimate of the model’s performance in the most frequent HLA-DP allele pairs.

| HLA-DP<br>Metric | A1*01:03<br>B1*02:01 | A1*01:03<br>B1*06:01 | A1*02:01<br>B1*05:01 | A1*02:01<br>B1*14:01 | A1*03:01<br>B1*04:02 |
| --- | --- | --- | --- | --- | --- |
| **AUC** | 0.94 | 0.95 | 0.80 | 0.73 | 0.78 |
| **F1** | 0.63 | 0.74 | 0.55 | 0.44 | 0.54 |
| **MCC** | 0.60 | 0.71 | 0.49 | 0.36 | 0.40 |

### Supplementary figures

**Supplementary Figure 1:**
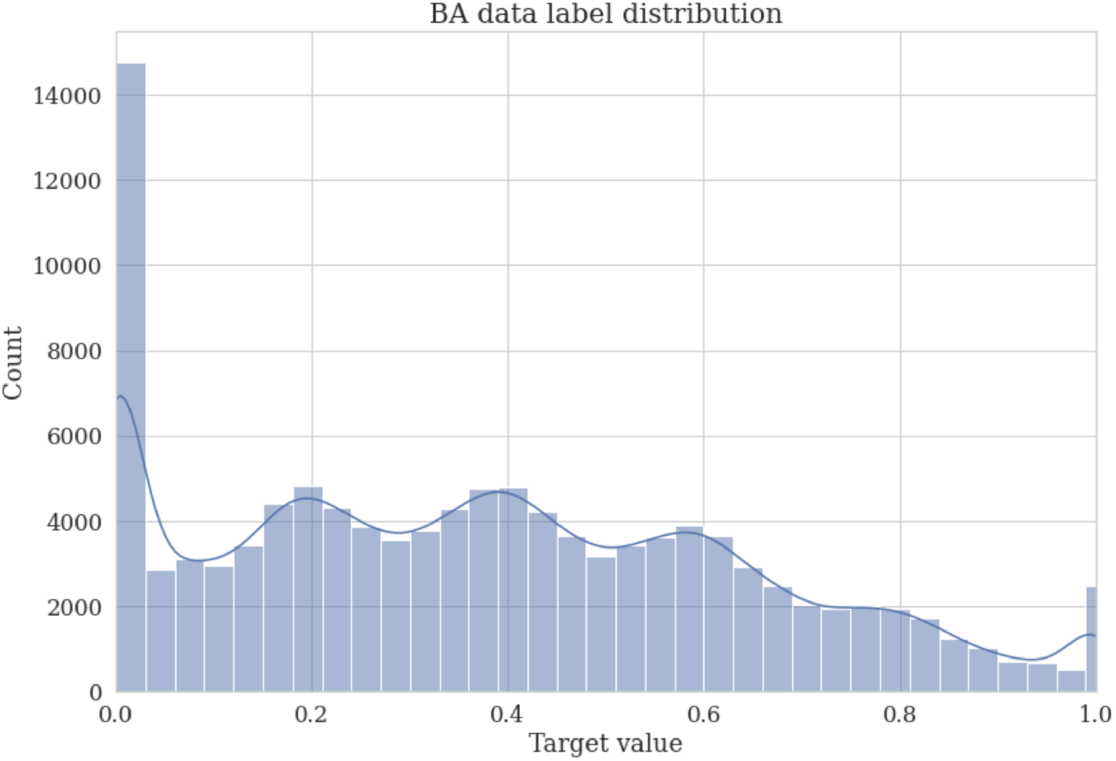
Histogram of label distributions of the BA data

**Supplementary Figure 2:**
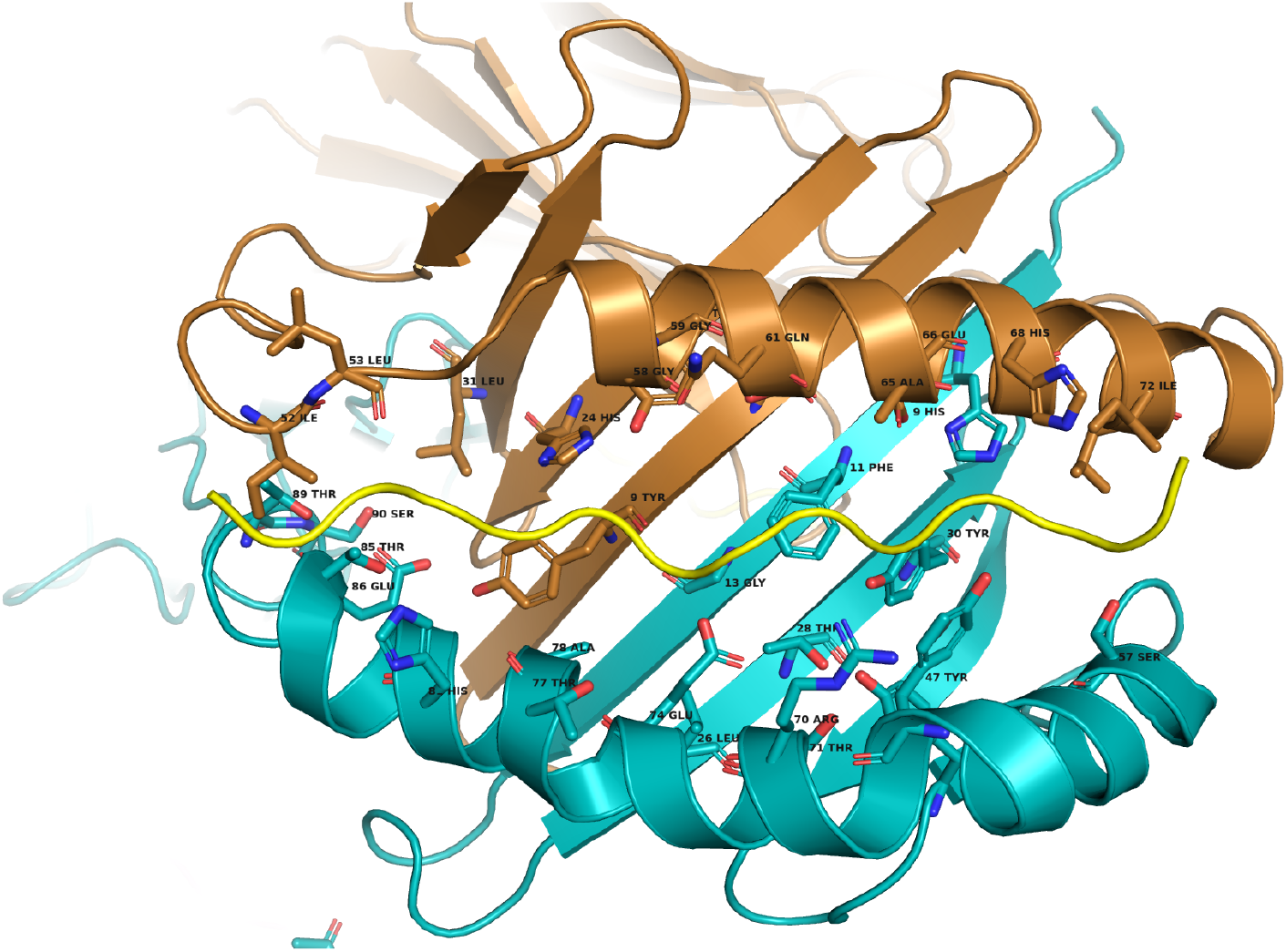
Cartoon representation of the I-A^g7^ structure (PDB code 1f3j). The Aa chain is colored in copper, the Ab1 chain in teal, and the binding peptide in yellow. The residues conforming the pseudo sequence are indicated using a *ball and stick* representation and labeled.

**Supplementary Figure 3:**
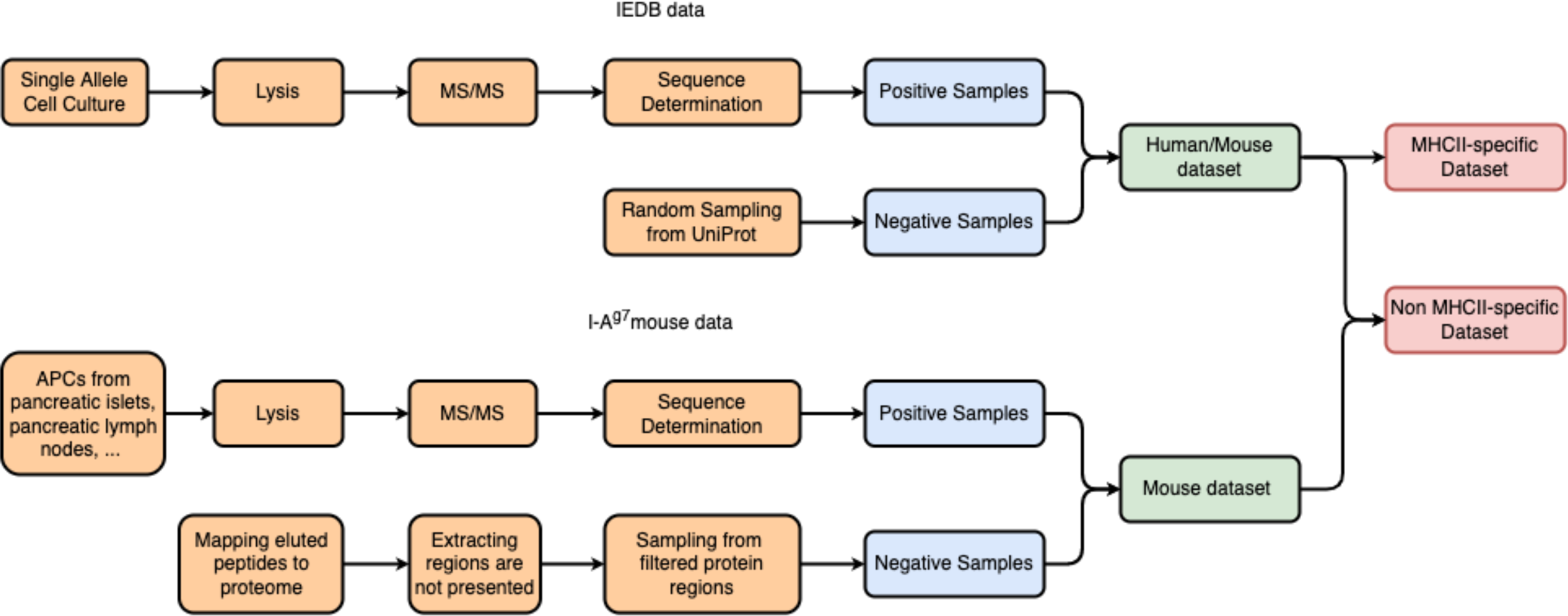
Overview of the data processing and generation workflow.

**Supplementary Figure 4:**
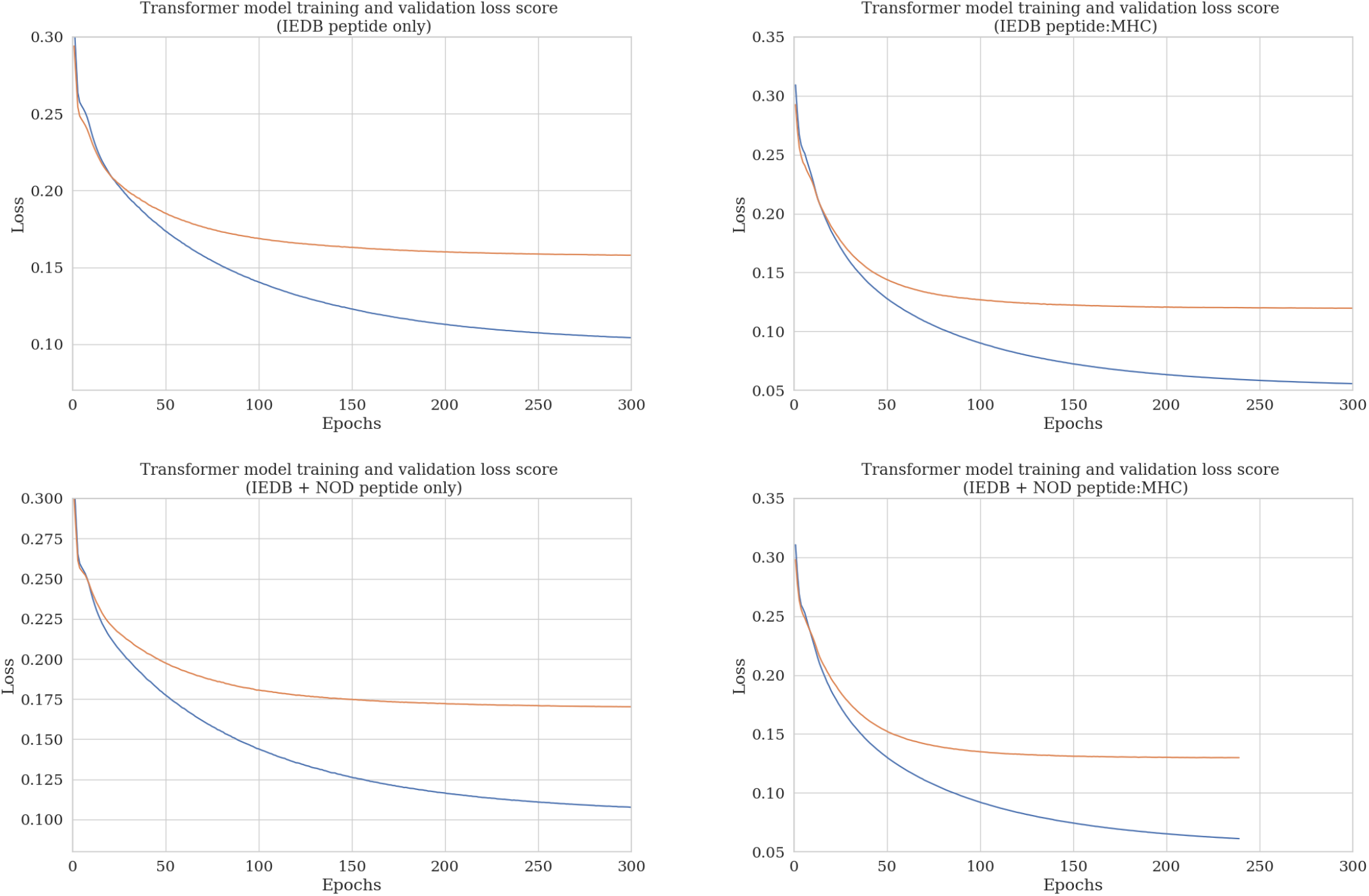
Transformer learning curves with confidence intervals from four different training runs. Data sources and feature sets are indicated in brackets.

## Notes

http://doi.org/10.5281/zenodo.7247911

### Summary of Updates

Table 1 Table 2 Table 3 Figure 2 Supplementary tables 1-4 Supplementary figure 4 The corresponding information in Results and Discussion has been consequently updated.

## References

Abelin, J. G., Keskin, D. B., Sarkizova, S., Hartigan, C. R., Zhang, W., Sidney, J., Stevens, J., Lane, W., Zhang, G. L., Eisenhaure, T. M., et al. (2017). Mass spectrometry profiling of HLA-associated peptidomes in mono-allelic cells enables more accurate epitope prediction. Immunity, 46(2):315–326.

Barra, C., Alvarez, B., Paul, S., Sette, A., Peters, B., Andreatta, M., Buus, S., and Nielsen, M. (2018). Footprints of antigen processing boost MHC class II natural ligand predictions. Genome medicine, 10(1):1–15.

Chen, B., Khodadoust, M. S., Olsson, N., Wagar, L. E., Fast, E., Liu, C. L., Muftuoglu, Y., Sworder, B. J., Diehn, M., Levy, R., et al. (2019). Predicting hla class ii antigen presentation through integrated deep learning. Nature biotechnology, 37(11):1332–1343.

Cheng, J., Bendjama, K., Rittner, K., and Malone, B. (2021). Bertmhc: improved mhc– peptide class ii interaction prediction with transformer and multiple instance learning. Bioinformatics, 37(22):4172–4179.

Devlin, J., Chang, M.-W., Lee, K., and Toutanova, K. (2019). Bert: Pre-training of deep bidirectional transformers for language understanding. In Proceedings of NAACL-HLT, volume 1, pages 4171–4186. Association for Computational Linguistics.

Graham, D. B., Luo, C., O’Connell, D. J., Lefkovith, A., Brown, E. M., Yassour, M., Varma, M., Abelin, J. G., Conway, K. L., Jasso, G. J., et al. (2018). Antigen discovery and specification of immunodominance hierarchies for mhcii-restricted epitopes. Nature medicine, 24(11):1762–1772.

Graham, D. B. and Xavier, R. J. (2020). Pathway paradigms revealed from the genetics of inflammatory bowel disease. Nature, 578(7796):527–539.

Greenbaum, J., Sidney, J., Chung, J., Brander, C., Peters, B., and Sette, A. (2011). Functional classification of class ii human leukocyte antigen (hla) molecules reveals seven different supertypes and a surprising degree of repertoire sharing across supertypes. Immunogenetics, 63:325–35.

Jarnuczak, A. F., Lee, D. C., Lawless, C., Holman, S. W., Eyers, C. E., and Hubbard, S. J. (2016). Analysis of intrinsic peptide detectability via integrated label-free and SRM-based absolute quantitative proteomics. Journal of proteome research, 15(9):2945–2959.

Jensen, K. K., Andreatta, M., Marcatili, P., Buus, S., Greenbaum, J. A., Yan, Z., Sette, A., Peters, B., and Nielsen, M. (2018). Improved methods for predicting peptide binding affinity to mhc class ii molecules. Immunology, 154(3):394–406.

Karle, A. C. (2020). Applying MAPPs assays to assess drug immunogenicity. Frontiers in immunology, 11:698.

Karosiene, E., Rasmussen, M., Blicher, T., Lund, O., Buus, S., and Nielsen, M. (2013). NetMHCIIpan-3. 0, a common pan-specific MHC class II prediction method including all three human MHC class II isotypes, HLA-DR, HLA-DP and HLA-DQ. Immunogenetics, 65(10):711–724.

Kingma, D. P. and Ba, J. (2014). Adam: A method for stochastic optimization. arXiv preprint arXiv:1412.6980.

Laimer, J. and Lackner, P. (2021). MHCII3D—Robust Structure Based Prediction of MHC II Binding Peptides. International journal of molecular sciences, 22(1):12.

Mueller, D. L. (2010). Mechanisms maintaining peripheral tolerance. Nature immunology, 11(1):21–27.

Murphy, T. L. and Murphy, K. M. (2021). Dendritic cells in cancer immunology. Cellular & Molecular Immunology, pages 1–11.

Nielsen, M. and Lund, O. (2009). Nn-align. an artificial neural network-based alignment algorithm for mhc class ii peptide binding prediction. BMC bioinformatics, 10(1):1–10.

Nielsen, M., Lundegaard, C., Blicher, T., Peters, B., Sette, A., Justesen, S., Buus, S., and Lund, O. (2008). Quantitative predictions of peptide binding to any HLA-DR molecule of known sequence: NetMHCIIpan. PLoS computational biology, 4(7):e1000107.

Nielsen, M., Lundegaard, C., Worning, P., Lauemøller, S. L., Lamberth, K., Buus, S., Brunak, S., and Lund, O. (2003). Reliable prediction of t-cell epitopes using neural networks with novel sequence representations. Protein Sci, 12(5):1007–17.

Pedregosa, F., Varoquaux, G., Gramfort, A., Michel, V., Thirion, B., Grisel, O., Blondel, M., Prettenhofer, P., Weiss, R., Dubourg, V., Vanderplas, J., Passos, A., Cournapeau, D., Brucher, M., Perrot, M., and Duchesnay, E. (2011). Scikit-learn: Machine learning in Python. Journal of Machine Learning Research, 12:2825–2830.

Pepper, M. and Jenkins, M. K. (2011). Origins of CD4+ effector and central memory T cells. Nature immunology, 12(6):467–471.

Racle, J., Michaux, J., Rockinger, G. A., Arnaud, M., Bobisse, S., Chong, C., Guillaume, P., Coukos, G., Harari, A., Jandus, C., et al. (2019). Robust prediction of HLA class II epitopes by deep motif deconvolution of immunopeptidomes. Nature biotechnology, 37(11):1283–1286.

Rajapakse, M., Lan Zhang, G., Srinivasan, K. N., Schmidt, B., Petrovsky, N., and Brusic, V. (2006). Prednod, a prediction server for peptide binding to the h-2g7 haplotype of the non-obese diabetic mouse. Autoimmunity, 39(8):645–650.

Reinherz, E. L., Tan, K., Tang, L., Kern, P., Liu, J.-h., Xiong, Y., Hussey, R. E., Smolyar, A., Hare, B., Zhang, R., et al. (1999). The crystal structure of a T cell receptor in complex with peptide and MHC class II. Science, 286(5446):1913–1921.

Reynisson, B., Alvarez, B., Paul, S., Peters, B., and Nielsen, M. (2020). NetMHCpan-4.1 and NetMHCIIpan-4.0: improved predictions of MHC antigen presentation by concurrent motif deconvolution and integration of MS MHC eluted ligand data. Nucleic acids research, 48(W1):W449–W454.

Schneidman-Duhovny, D., Khuri, N., Dong, G. Q., Winter, M. B., Shifrut, E., Friedman, N., Craik, C. S., Pratt, K. P., Paz, P., Aswad, F., et al. (2018). Predicting CD4 T-cell epitopes based on antigen cleavage, MHCII presentation, and TCR recognition. PloS one, 13(11):e0206654.

Stern, L. J. and Santambrogio, L. (2016). The melting pot of the MHC II peptidome. Current opinion in immunology, 40:70–77.

Ten Broeke, T., Wubbolts, R., and Stoorvogel, W. (2013). MHC class II antigen presentation by dendritic cells regulated through endosomal sorting. Cold Spring Harbor perspectives in biology, 5(12):a016873.

UniProt Consortium (2021). UniProt: the universal protein knowledgebase in 2021. Nucleic acids research, 49(D1):D480–D489.

Vaswani, A., Shazeer, N., Parmar, N., Uszkoreit, J., Jones, L., Gomez, A. N., Kaiser, £., and Polosukhin, I. (2017). Attention is all you need. In Advances in neural information processing systems, pages 5998–6008.

Vita, R., Mahajan, S., Overton, J. A., Dhanda, S. K., Martini, S., Cantrell, J. R., Wheeler, D. K., Sette, A., and Peters, B. (2019). The immune epitope database (IEDB): 2018 update. Nucleic acids research, 47(D1):D339–D343.

Wan, X., Vomund, A. N., Peterson, O. J., Chervonsky, A. V., Lichti, C. F., and Unanue, E. R. (2020). The MHC-II peptidome of pancreatic islets identifies key features of autoimmune peptides. Nature immunology, 21(4):455–463.

Wieczorek, M., Abualrous, E. T., Sticht, J., Álvaro-Benito, M., Stolzenberg, S., Noé, F., and Freund, C. (2017). Major histocompatibility complex (MHC) class I and MHC class II proteins: conformational plasticity in antigen presentation. Frontiers in immunology, 8:292.

Wu, Y., Zhang, N., Hashimoto, K., Xia, C., and Dijkstra, J. M. (2021). Structural comparison between mhc classes i and ii; in evolution, a class-ii-like molecule probably came first. Front Immunol, 12:621153.

Zhang, Z., Luo, L., Xing, C., Chen, Y., Xu, P., Li, M., Zeng, L., Li, C., Ghosh, S., Della Manna, D., et al. (2021). Rnf2 ablation reprograms the tumor-immune microenvironment and stimulates durable nk and cd4+ t-cell-dependent antitumor immunity. Nature Cancer, 2(10):1018–1038.

